# E484K and N501Y SARS-CoV 2 Spike Mutants Increase ACE2 Recognition but Reduce Affinity for Neutralizing Antibody

**DOI:** 10.1101/2021.06.23.449627

**Authors:** Sandipan Chakraborty

**Author notes:** Corresponding author: Sandipan Chakraborty.

## Abstract

SARS-CoV2 mutants emerge as variants of concern (VOC) due to altered selection pressure and rapid replication kinetics. Among them, lineages B.1.1.7, B.1.351, and P.1 contain a key mutation N501Y. B.1.135 and P.1 lineages have another mutation, E484K. Here, we decode the effect of these two mutations on the host receptor, ACE2, and neutralizing antibody (B38) recognition. The gain in binding affinity for the N501Y RBD mutant to the ACE2 is attributed to improved π-π stacking and π-cation interactions. The enhanced receptor affinity of the E484K mutant is caused due to the formation of a specific hydrogen bond and salt-bridge interaction with Glu75 of ACE2. Notably, both the mutations reduce the binding affinity for B38 due to the loss of several hydrogen-bonding interactions. The insights obtained from the study are crucial to interpret the increased transmissibility and reduction in the neutralization efficacy of rapidly emerging SARS-CoV2 VOCs.

## 1. Introduction

SARS-CoV2 emerged initially from a local seafood market in Wuhan, China, is now a pandemic that causes severe outbreaks in more than 216 countries. As per the data on 14^th^ June 2021, the total infected cases are 17.6 crores with 38.1 lacs mortality. The disease outbreak started in December 2019. After that, the virus evolved during the pandemic. Genetic diversity of the host population drives the SARS-CoV2 genomic variations due to altered selection pressure.^1^ High mutational rates and rapid replicative kinetics lead to the diversification of SARS-CoV2 genomes.

SARS-CoV2 is a positive-stranded RNA virus enclosed within a viral envelope.^2^ Three proteins, E, M, and spike glycoprotein, are embedded in the viral envelope.^2^ Among them, spike protein forms a large clover-shaped protrusion as a homo-trimer recognizing the human ACE2 receptor to mediate viral entry.^3^ Spike monomer consists of the S1 and S2 domains. Three S1 domains from the three trimers associate to form the ectodomain, and the S2 domains entangle to create the stalk, transmembrane, and small intracellular domains.^2,4^ The receptor-binding domain (RBD) of the S1 binds to the peptidase domain (PD) of the ACE2 receptor to open up the S2 cleavage site. The cleavage by the host proteases mediates the fusion of the viral membrane to the host membrane.^4,5^ Due to its role in host receptor recognition, spike protein is under positive selection pressure to produce SARS-CoV2 variants with increased transmissibility and infection rate.^6^ Spike protein is the target for vaccine and immunogenic therapy development as there exist many immunodominant solvent-exposed epitopes that are readily accessible by antibody pool.^7^ Thus, tracking the SARS-CoV2 spike variant is very important to identify mutant viral strains with higher transmissibility and the ability to cause immune invasion.

D614G is the first fitness-enhancing spike mutation, which became the significant circulating strain from April 2020 onwards. Calculation of the dN/dS ratio suggests a positive selection bias at the 614^th^ codon position with increased infectivity.^8^ Data indicate that the viral genome acquires ~2-3 mutations per month.^9^ Although most of the mutations purge out from the population, few of them are fitness-enhancing mutations that alter the antigenic phenotype of SARS-CoV2, which require focused attention. Previously, our group reported two spike mutants, V367F and S494P, that showed enhanced human ACE2 binding ability.^10^ Later, it was demonstrated that the S494P caused a 3-5 fold decrease in neutralization titer^11^ and was reported in many UK, USA, and Mexico cases.^12^ Among several identified spike mutants, mutations that occur within the RBM (Receptor Binding Motif; residues 438◻506) deserve special attention. They can increase human ACE2 affinity as well as decrease neutralization by several mABs. To date, many significant spike RBD mutants were identified with altered ACE2 recognition and antigenic properties. N439K, L452R, and Y453F showed an increase in ACE2 receptor binding ability, whereas G446V, S477N, G485R, and F490S demonstrated ~3-5 fold decrease in neutralization titer for a few sera.^13^ WHO declared several SARS-CoV2 mutants as variants of concern (VOC) as they cause sustained disease outbreak across several regions of the globe. Among them, lineages B.1.1.7, B.1.351, and P.1 contain a key common RBM mutation, N501Y, which was experimentally shown to increase the ACE2 affinity.^14^ In pseudo-viruses carrying the N501Y mutation, a 10-fold decrease in efficacy was reported during the neutralization of mRNA vaccine◻elicited mAbs.^15^ However, mouse◻adapted SARS◻CoV◻2 N501Y strain can be effectively neutralized by vaccine◻elicited sera.^16^ Two other rapidly emerging VOCs (B.1.135 and P.1) contain another crucial RBM mutation, E484K. Also, the variant of interest P.2, first reported in Rio de Janeiro and then rapidly widespread in the northeast region of Brazil, contains only the E484K spike mutation.^13^ This mutation may be accountable for evasion from neutralizing antibodies.^17,18^ Recently, *in-vitro* micro-neutralization assays revealed a significant reduction in neutralization efficiency for the recombinant (r)SARS-CoV-2 virus with E484K mutation compared to the control USA-WA1/2020 strain on 34 sera collected from different study participants.^18^ Also, it was demonstrated that the E484K variant caused a 3·4-fold decrease in the neutralization titer in five individuals who received two doses of the Pfizer–BioNTech vaccine.^18^

Thus, both the N501Y and E484K mutations appear to play critical roles in the emergence of many VOCs with higher infectivity rates and immune invasion ability. But the mechanism is yet to be understood at the molecular level, which is highly crucial to understand the efficacy of mutant strains on neutralization sera and vaccines. Here, using extensive all-atom molecular dynamics simulation and free energy calculations, we critically decode the role of both the N501Y and E484K spike mutations on ACE2 and neutralization antibody recognition.

## 2. Materials and Method

### 2.1. Preparation of spike mutants, spike-ACE2, and spike-antibody complexes

Recently, I refined the crystal structure of the SARS-CoV2 RBD-ACE2 complex (PDB ID: 6M0J^19^) using extensive molecular dynamics simulation, which was used as a starting structure for the present study.^10^ The E484K and N501Y mutant complexes were built from the wild-type SARS-CoV2 RBD-ACE2 complex by mutating the glutamate to Lysine and asparagine to tyrosine, respectively, in the spike RBD region.

The recently resolved crystal structure of the SARS-CoV2 spike RBD complexed with a neutralizing antibody (B38) was considered for the study (PDB ID:7BZ5).^20^ The Wild-type SARS-CoV2 RBD from the RBD-ACE2 complex was aligned on the RBD-B38 crystal structure, and then the aligned RBD with the B38 antibody was considered as the wild-type RBD-B38 docked complex. The E484K-B38 and N501Y-B38 complexes were prepared by carrying out the specific mutations on the wild-type SARS-CoV2 RBD-B38 complex using the mutagenesis toolkit of Visual Molecular Dynamics (VMD)^21^.

### 2.2. Equilibrium simulations of wild-type and mutant SARS-CoV2 spike RBD-ACE2 and RBD-antibody complexes

All the simulations were performed using GROMACS 2018.1^22,23^ packages using the AMBER99SB-ILDN force field.^24^ All the complexes were first energy minimized *in vacuo* to remove any bad contacts. Then each complex was immersed in a triclinic box so that the minimum distance between any protein atom and box walls was >10 ◻. The box dimensions for wild-type and mutant SARS-CoV2 RBD-ACE2 complexes were 100 × 100 × 180 ◻^3^, and for wild-type and mutant SARS-CoV2 RBD-B38 complexes, the box dimensions were 100 × 100 × 190 ◻^3^. Each box was solvated with TIP3P water, and an appropriate number of counter ions were added to neutralize the charge of each system. Then, 500 steps of energy minimization using the steepest descent algorithm were carried out for each system, followed by 10ns of position-restrained dynamics where the protein backbone dynamics were restrained. At the same time, water molecules were allowed to move freely. After that, a 2ns NVT simulation was carried out for each complex at 298 K, followed by another 2ns NPT simulation where both the proteins and solvent molecules were allowed to move freely. Finally, 500 ns of production simulations were performed in the NPT ensemble. All the simulations were carried out under periodic boundary conditions. The temperature was kept constant by coupling to a Nosé–Hoover thermostat with a coupling time constant of 0.1 ps. The pressure was maintained at 1 bar thorough coupling to the isotropic Parrinello–Rahman barostat with the time constant for coupling set to 2 ps. Electrostatic interactions were calculated using the PME method with default values for grid spacing.

### 2.3. Calculation of the potential of mean force (PMF) for wild-type and mutant spike RBD to ACE2 and B38 antibody

The optimized wild-type and mutant (E484K and N501Y) SARS-CoV2 RBD complexed with ACE2 and B38 were immersed in a triclinic box filled with TIP3P water such that the minimum distance between any protein atom and box edges was > 10 ◻. The box dimensions for wild-type and mutant SARS-CoV2 RBD-ACE2 complexes were 100 × 100 × 200 ◻^3^, and for wild-type and mutant SARS-CoV2 RBD-B38 complexes, the box dimensions were 100 × 100 × 190 ◻^3^. In Z-direction, the box length was chosen in a way that it was greater than the double of final pull distance. Charges of each system were neutralized by adding an appropriate number of counterions. Each system was then minimized with 500 steps of the steepest descent algorithm. Then, 10ns position restrained dynamics were performed where the proteins were restrained while water molecules were allowed to move freely. This was followed by 2ns equilibration at 298 K in the NVT ensemble, and 2ns NPT simulation performed using the same simulation protocol mentioned in the equilibrium simulation section.

The potential of mean force (PMF) for pulling the spike RBD (Wild-type and mutants) from the ACE2 or B38 was computed using the umbrella sampling techniques. RBD was pulled from the ACE2/B38 protein binding interface along the Z-direction with an interval of 1 ◻ with an umbrella force constant of 500 kJ.mol^−1^.nm^2^. In each umbrella window, 2ns equilibration was performed, followed by 3ns production run in NPT ensemble using the same thermostat, barostat, and associated coupling parameters, mentioned above. Two four windows were considered for each case to sample the entire reaction coordinate. A weighted histogram analysis method (WHAM)^25^ was used to construct the PMF profile. Sufficient overlap among all the windows was confirmed by histogram analysis. For each system, at least three independent umbrella simulations were performed.

## 3. Result and Discussions

### 3.1. Interactions of wild-type, N501Y and E484K mutant RBDs with ACE2: Equilibrium simulation and binding free energy analysis

The RBD structure was built using the sequence of SARS-CoV2 samples collected from the Wuhan seafood market (isolate=Wuhan-Hu-1), referred to as wild-type in the text. Equilibrium molecular dynamics simulations have been used to underscore the effect of the E484K and N501Y RBD mutations on ACE2 recognition. A root mean square deviation (RMSD) based clustering of the simulation trajectories for the three systems (Wild-type RBD-ACE2, N501Y RBD-ACE2, and E484K-ACE2) is shown in Fig. 1A. Upon complexed with ACE2, the wild-type RBD is comparatively stable with two evident conformational clusters. The larger cluster contains RBD conformations observed during the 120-500 ns simulation timescale.

**Fig. 1:**
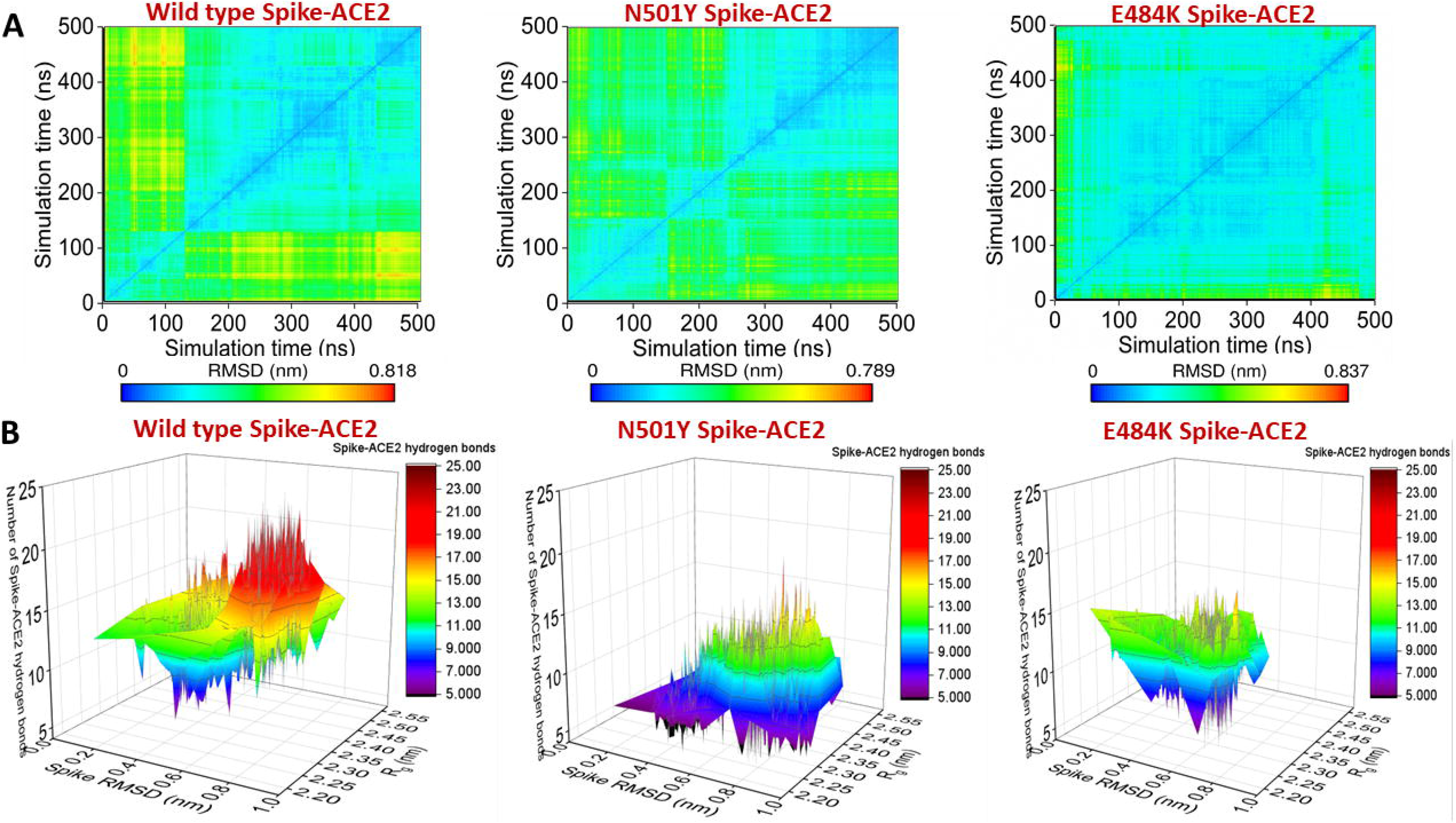
(A) Root mean square deviation (RMSD) based clustering of the SARS-CoV2 spike RBD-ACE2 complexes obtained from the simulation trajectories for the three systems (Wild-type RBD-ACE2, N501Y RBD-ACE2, and E484K-ACE2) is shown. (B) 3-D representation of the conformational dynamics obtained from the simulations of all the three complexes in terms of the root mean square deviation (RMSD), the radius of gyration (Rg) space of RBD, and RBD-ACE2 hydrogen bonds.

The N501Y mutation in spike RBM induces dynamics in the complex. Three different conformational clusters are evident. Conformations from the first 120 ns of the simulation are clubbed together in a cluster. The second cluster contains conformations observed during the 120-200 ns timescale. All the conformations from the last 300 ns clubbed into the third cluster. On the other hand, the E484K RBD mutation significantly stabilizes the complex. All the conformations are clubbed into a single cluster.

3-D representation of the conformational dynamics obtained from the simulations of all the three complexes is shown in Fig. 1B. In the root mean square deviation (RMSD) and radius of gyration (R_g_) space, wild-type RBD dynamics are confined rather narrowly compared to the N501Y mutant when complexed with the ACE2. In this conformational ensemble, the RBD-ACE2 hydrogen bonding varies greatly. On average, there are 11-13 RBD-ACE2 hydrogen bonds. Two populations are evident, one with 7-9 RBD-ACE2 hydrogen bonds and the other one with 17-19 RBD-ACE2 hydrogen bonds. In contrast, N501Y mutant RBD forms less number of interfacial hydrogen bonds. A small proportion of the complex conformations are observed with 7-9 hydrogen bonds, while the rest of the complex conformations are characterized by ~11-13 RBD-ACE2 hydrogen bonds. The conformational space sampled during the molecular dynamics simulations by the E484K spike mutant is more confined in RMSD and R_g_ space, reconfirming high stabilization of the complex. The average number of interfacial hydrogen bonds ranges between 13 to 15 for most of the sampled conformations. A small conformational state with reduced RBD-ACE2 hydrogen bonds (7-9 hydrogen bonds) is also noticed.

Further RMSD based clustering is used to identify the most populated solution structure of wild-type, N501Y, and E484K RBD complexed with ACE2. The wild-type complex visited 151 conformational clusters during the simulation using a 1.2 ◻ RMSD cut-off. The 136^th^ cluster is the most populated, span over 120-500 ns during the simulation (Fig. 2A). N501Y mutation increases the number of clusters, and a total of 180 clusters are observed (Fig. 2A). Many small conformational clusters are evident from the simulation trajectory. Among them, the 178^th^ cluster is the most populated one.

**Fig. 2:**
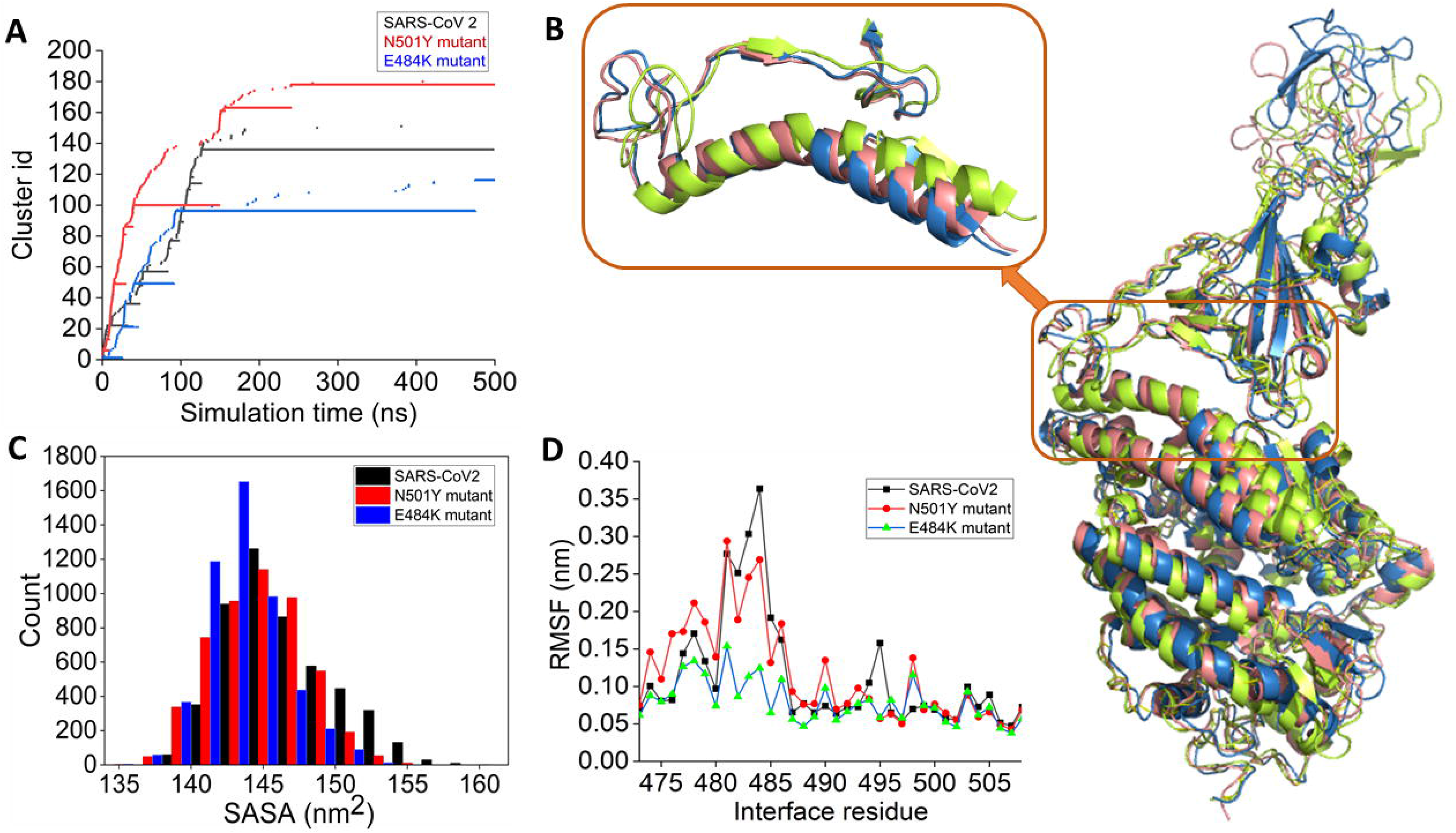
(A) Time-evolution of conformational clusters evident from the RMSD based clustering of simulation trajectory of wild-type (black), N501Y (red), and E484K (blue) RBD complexed with ACE2. (B) The alignment of the average complex structure from the most populated clusters for the three systems is shown. Wild-type, N501Y, and E484K RBD complexed with ACE2 are colored as deep salmon, greenish-yellow, and blue, respectively. (C) Distribution of the solvent-accessible surface area (SASA) of wild-type and mutant RBDs obtained from the simulations of three RBD-ACE2 complexes. (D) Root mean square fluctuations (RMSF) of the receptor binding motif obtained from the simulations of RBD-ACE2 complexes.

E484K RBD-ACE2 complex visited the least number of clusters during the simulation, further indicating high stabilization of the complex. The 96^th^ cluster is the most populated one. The alignment of the average complex structure from the most populated clusters for the three complexes is shown in Fig. 2B. Overall the RBD-ACE2 complex structures remain very similar, apart from the fluctuations of several loop regions. The binding interface has been zoomed in the inset. The wild-type and E484K RBD-ACE2 complexes show similar interfacial packing. However, the N501Y RBD mutation alters interfacial packing. Notably, the N-terminal loopy overhang region that packs the interfacial helical region of the peptidase domain of the ACE2 changes its conformation such that it loses contacts with the ACE2.

Analysis of the solvent-accessible surface area (SASA) of RBD reveals that the interfacial area packed between the mutants (N501Y and E484K) spike RBD and ACE2 are higher than the wild-type RBD. The E484K RBD is packed tightly to the ACE2 PD surface during the simulation (Fig. 2C). Root mean square fluctuations also reveal that E484K mutation highly stabilized the RBM upon complexation with the ACE2. The region from 480-486 is highly stabilized in E484K mutated RBD compared to the wild-type. N501Y mutation in the RBD also mildly stabilizes this region (Fig. 2D).

Equilibrium simulations indicate that both N501Y and E484K mutations stabilize the RBM upon complexation with ACE2. To evaluate the effect of stabilization of mutant RBD upon binding leads to an increase in the binding affinity for ACE2, the potential of mean force (PMF) has been computed. The free energy for complex formation (∆G_bind_) is obtained by calculating the unbinding energy of wild-type and mutant RBDs from the RBD-ACE2 complexes using the umbrella sampling method. Wild-type RBD binds to the ACE2 with an affinity of −144 kJ/mol. The N501Y mutant RBD interacts more strongly with the ACE2. The calculated binding free energy is −165 kJ/mol. The E484K mutant RBD showed a remarkably higher affinity towards ACE2 with the calculated binding free energy of −210 kJ/mol (Fig. 3A). We then decode the gain of the ACE2 binding energy for mutant RBDs in terms of interactive features. Fig 3B represents the colored coded representation of interfacial interaction involved in ACE2 recognition by spike RBDs for all three cases, and Fig. 3C, 3D, 3E, and 3F show the mapping of those interactions on the RBD-ACE2 interface for wild-type and mutant spike proteins. Recently, Chakraborty *et al.* showed that the Phe486 is the highest energetic contributor for forming the spike-ACE2 complex using the MM/GBSA method.^10^ This interaction is also preserved for mutants too. The residue interacts with the Met82 of ACE2 using van der Waals interactions. TYR489 is another significant energetic contributor that interacts with Phe486. This interaction is evident in the wild-type and N501Y mutant but lost upon E484K mutation. The E484K mutation disrupts most of the hydrophobic interactions involved in ACE2 recognition. Upon N501Y mutation, the Tyr501 gains van der Waals contacts with the ACE2 interface. Fig. 3C represents the common interfacial hydrogen bonds evident in all the three RBD-ACE2 complexes. Hydrogen bonding interactions involving the Gln493, Arg403, Gly496, Gln498, and Thr500 present at the middle and C-terminal loop of the RBM preserve in all the three RBD-ACE2 complexes.

**Fig. 3:**
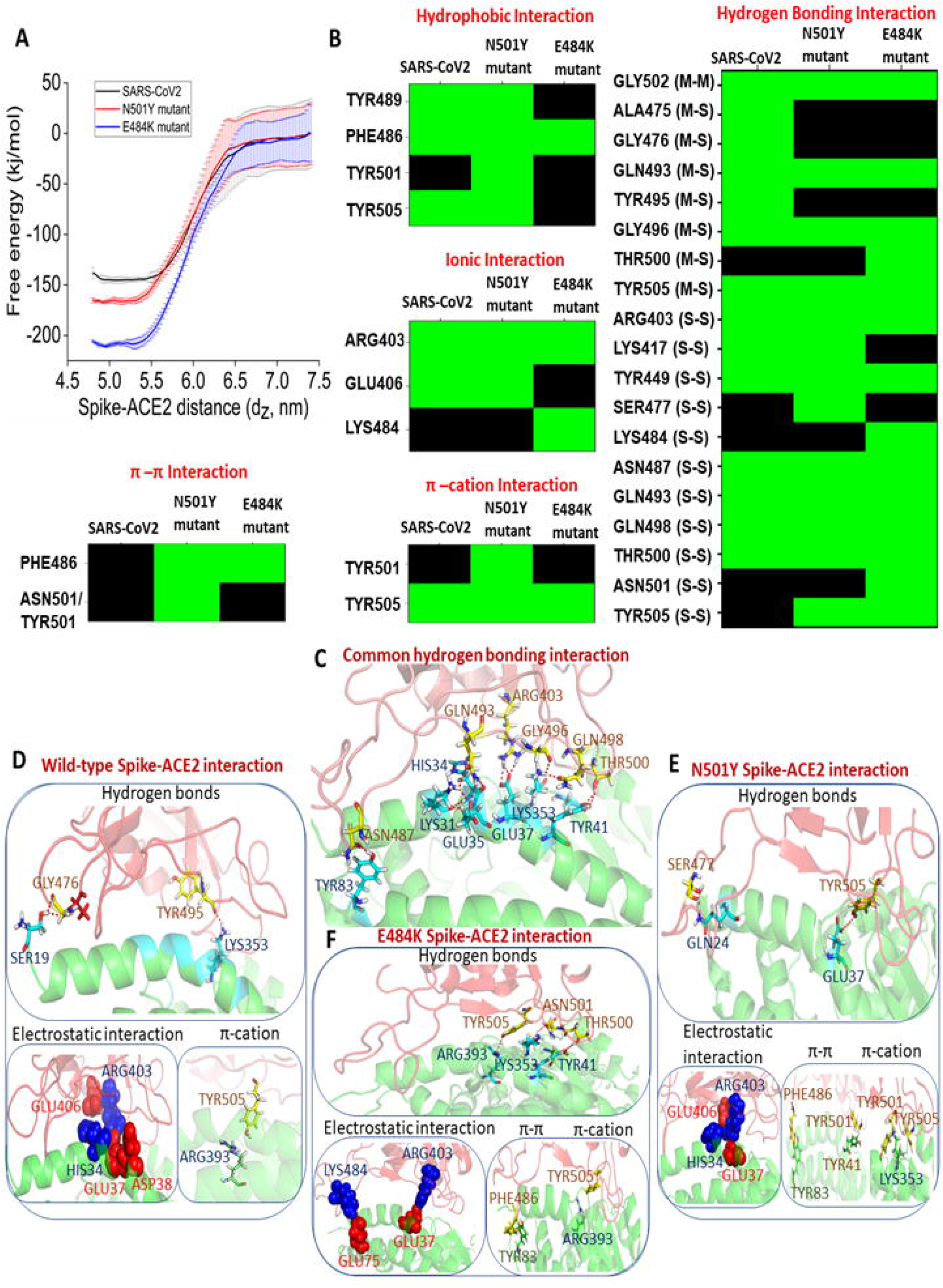
(A) Potential of mean force for the binding of wild-type (black), N501Y (red), and E484K (blue) RBD to ACE2. (B) Color-coded representation of interfacial interactions involved in ACE2 recognition by the wild-type and mutant spike RBDs. (C) Display of common hydrogen-bonding interactions present in all the three RBD-ACE2 complexes. Protein is rendered in cartoon mode and residues forming hydrogen bonding interactions are shown in stick mode. Unique interactions present in the wild-type RBD-ACE2 complex (D), N501Y RBD-ACE2 complex (E), and E484K RBD-ACE2 complex (F) are shown. RBD and ACE2 are colored in red and green, respectively. Residues forming hydrogen bonds, π-π, and π-cation interactions are shown in stick mode. Residues involved in electrostatic interactions are shown in the sphere representation. Positive and negatively charged residues are colored in blue and red, respectively.

In addition, Wild-type RBD forms three unique hydrogen bonds between the Ala475, Gly476, and Tyr495 of the RBM and Ser19, Lys353 of ACE2 (Fig. 3D), which are absent in both the mutants. N501Y mutant RBD forms a specific hydrogen bond involving Ser477 of the RBD. The gain in binding affinity for the N501Y mutant to the ACE2 is due to the improved π-π and π-cation interactions. Mutation of asparagine to tyrosine at the 501^st^ position allows formation for a π-cation interaction with the Lys353 and a π-π stacking interaction with Tyr41 of ACE2 (Fig. 3E).

The E484K mutation allows Lys484 of the RBM to form specific hydrogen bonds with ACE2. In addition, the particular residue is involved in salt-bridge interaction with Glu75 of ACE2. These high-affinity interactions allow E484K mutant RBD to be firmly bound to the ACE2 interface. This intense encounter of E484K RBD to ACE2 allows the remodeling of few interfacial residues. The Tyr505 forms hydrogen bonding interactions with Arg393, Asn501 forms interactions with Lys353, and the Thr500 also form hydrogen-bonding interactions with Tyr41 of ACE2 (Fig. 3F). The Phe486 of RBD creates additional π-π stacking interaction with Tyr83 of ACE2, which is also present in the N501Y RBD-ACE2 complex.

The work highlights that the E484K and N501Y are gain-of-function mutants. By specific modulation of intermolecular hydrogen bonds, π-π stacking, and π-cation interactions, both the mutants bind to the host receptor with increased affinity.

### 3.2. Interactions of wild-type, N501Y and E484K mutant RBDs with a neutralizing antibody, B38: Equilibrium simulation and binding free energy analysis

Then, the effect of these two mutations on neutralizing antibody recognition has been evaluated using equilibrium simulation and free energy calculations. Recently, four human-origin monoclonal neutralizing antibodies (mAB) were identified from a convalescent patient.^20^ Among them, the B38 antibody binds to the spike RBD and competes with the ACE2. The crystal structure of the B38-RBD complex was also resolved at high resolution.^20^ The complex structure is used to explore the effect of both the E484K and N501Y RBM mutations on the B38 monoclonal antibody recognition.

In the 2-D space defined by RMSD and R_g_, the wild-type RBD-B38 complex occupies a distinct space compared to the two mutant-B38 complexes. As a result, the conformational ensemble sampled during the simulation timescale for the wild-type SARS-CoV2 RBD-B38, E484K RBD-B38, and N501Y RBD-B38 is different with almost no overlap (Fig. 4A). This altered conformational dynamics can result from differential interactions of mutant spikes with the heavy (H) and light (L) chains of the antibody.

Further, cluster analysis has been performed to identify the most populated solution structure of all the three RBD-antibody complexes. The 4^th^, 5^th^, and 1^st^ cluster conformations are almost stable for the entire simulation timescale for wild-type RBD-B38, E484K RBD-B38, and N501Y RBD-B38, respectively (Fig. 4B). Therefore, the representative conformation from each cluster has been identified for each complex, aligned, and shown in Fig. 4C. The loopy overhang of the RBM, which is in contact with the H and L chain of the B38 antibody, is highly stable without any noticeable deformation. However, the distal loop regions of the RBD show high flexibility. A closer look at the contact regions reveals the difference in the recognition mechanism (Fig. 4C, inset). The RBM forms many contacts with both the H and L chains, and E484K mutant RBM shows very similar protein-protein contacts with the binding interface of the B38 antibody. However, the RBM of the N501Y mutant spike loses critical contact with the H-chain of the antibody due to the upward curvature of a loop region. (Inset of the Fig. 4C). Wild-type SARS-CoV2 RBD forms a higher number of hydrogen bonds with the H-chain. Both the mutants show a significant reduction in the number of hydrogen bonds involving the H-chain of the antibody (Fig. 4D). Particularly, the E484K RBD loses a higher number of hydrogen bonding interactions during the simulation. The wild-type RBD forms a lower number of hydrogen bonding interactions with the L-chain than the H-chain. N501Y mutation further decreases the hydrogen bonding interaction with the L-chain. But the E484K mutant RBD shows an increase in hydrogen bonding interactions with the L-chain during the simulation. However, the distribution is broad with a reduction in the occurrence frequency, indicating the transient nature of these hydrogen-bonding interactions.

**Fig. 4:**
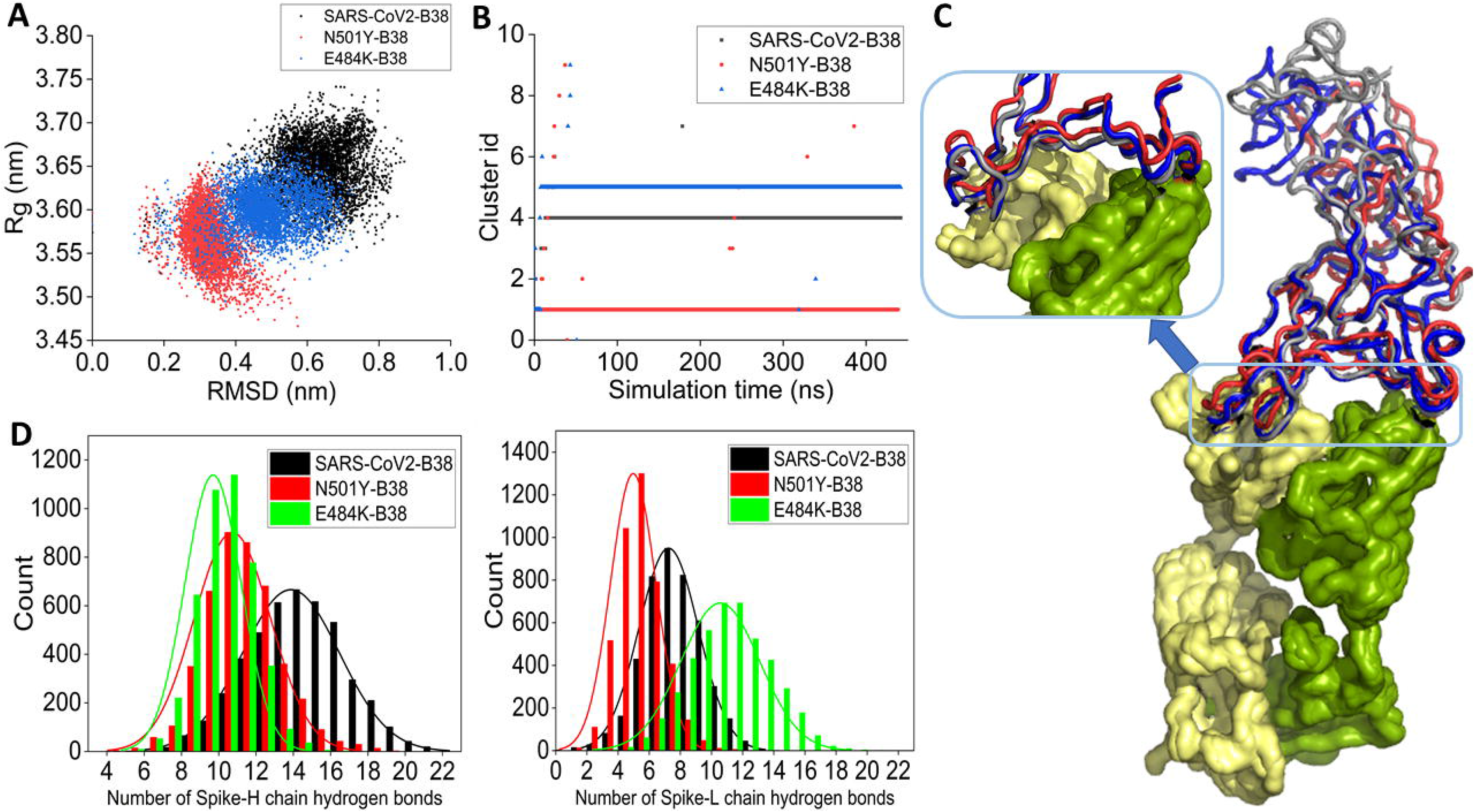
(A) 2-D scatter representation of the conformational ensemble of wild-type RBD-B38 (black), N501Y RBD-B38 (red), and E484K RBD-B38 (blue) complexes obtained from the equilibrium simulation in RMSD and R_g_ space. (B) Time-evolution of conformational clusters of wild-type (black), N501Y (red), and E484K (blue) RBD complexed with B38. (C) The alignment of the average complex structure from the most populated clusters for the three systems is shown. Wild-type, N501Y, and E484K RBD are colored gray, red, and blue. B38 is represented in surface mode. (D) The distribution of the number of hydrogen bonds between the wild-type and mutant RBDs and the H-chain (left) and L-chain (right) of the antibody.

Furthermore, the binding affinity of the wild-type and mutant RBDs for the antibody was calculated using umbrella sampling. Astonishingly, the potential of mean force (PMF) profiles reveals a reverse order of affinity compared to the ACE2 recognition. The wild-type RBD shows the highest affinity for the B38 mAB, and mutations reduce the binding affinity. Notably, the E484k mutant showed the least binding affinity to the B38 mAB (Fig. 5A).

**Fig 5:**
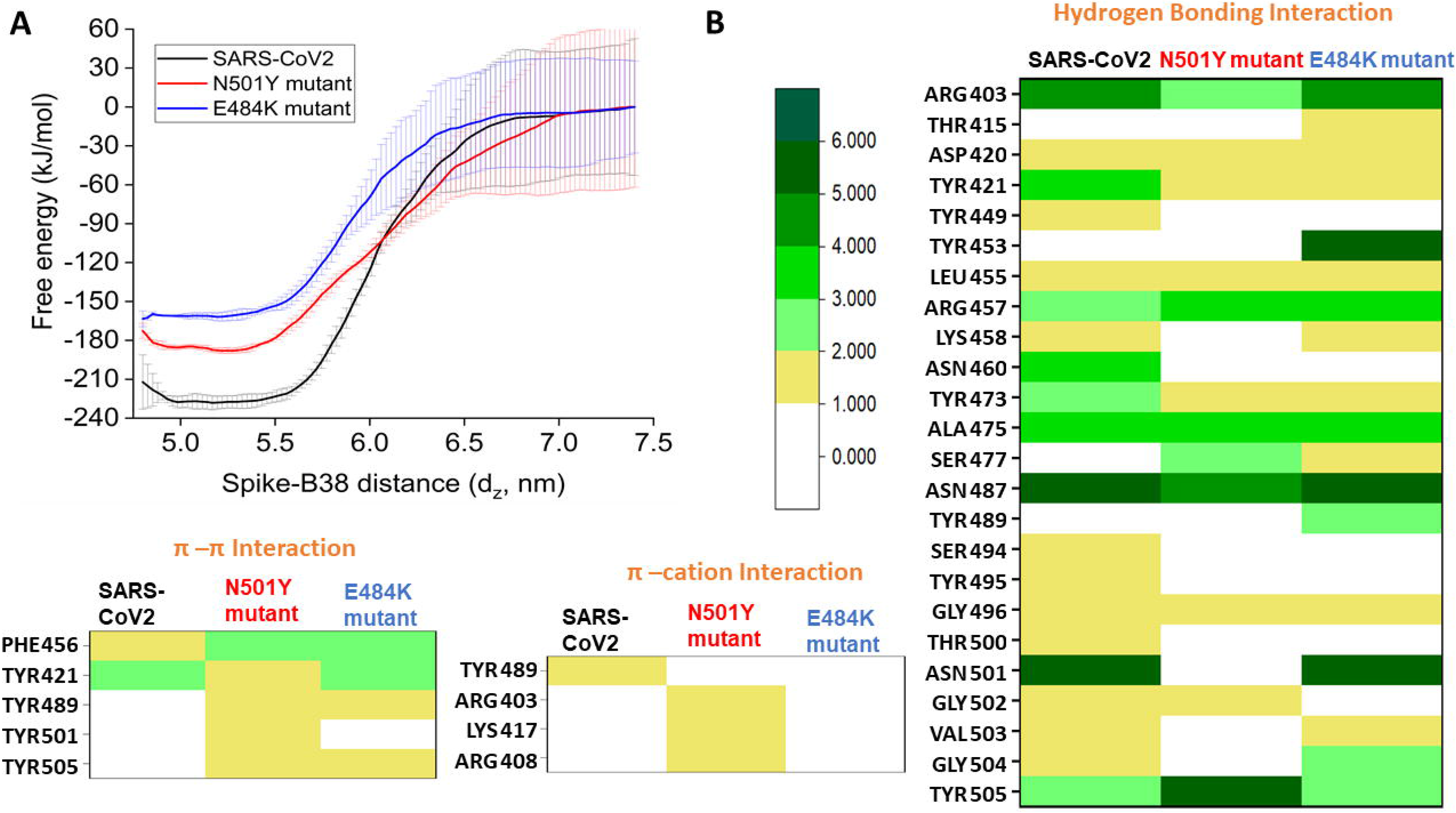
(A) Potential of mean force for the binding of wild-type (black), N501Y (red), and E484K (blue) RBD to B38 antibody. (B) Color-coded representation of interfacial interactions involved in the B38 antibody recognition by the wild-type and mutant spike RBDs. The frequency of interactions is scaled according to the color bar.

Then, the pharmacophoric feature for antibody recognition has been decoded for both the wild-type and mutant RBDs, and the results are shown in Fig. 5B. The N501Y mutant RDB gains few π-π stacking and π-cation interactions during antibody recognition. However, the loss of binding affinity is primarily due to the loss of crucial hydrogen bonding interactions. Asn501 in wild-type RBD forms six hydrogen bonds with the B38 antibody involving both the sidechain and mainchain donors and acceptors. Mutation of the residue with tyrosine leads to the complete loss of the entire hydrogen bonding network. Apart from this specific loss, Tyr449, Lys458, Asn460, Ser494, Tyr495, Thr500, Val503, and Gly504 also lose their hydrogen-bonding interactions with the B38 mAB. Thus, the loss of many hydrogen-bonding interactions with the antibody accounts for the reduced binding affinity of the N501Y mutant. The E484K mutant RBD loses all the π-cation interactions in addition to the loss of several hydrogen-bonding interactions with the B38 antibody. Tyr449, Asn460, Ser494, Tyr495, Thr500, and Gly502 lose hydrogen bonding interactions with the antibody present in the wild-type RBD-B38 complex.

## 4. Conclusion

The present study critically explores the effect of two critical mutations, E484K and N501Y, observed in few recently emerging variants of concern of SARS-CoV2. These two mutations occur at the receptor binding motif (RBM), which is involved in host receptor recognition as well as binds to the class I antibody. Equilibrium simulation and free energy calculations reveal both the mutations increase the affinity for ACE2 while showing a considerable reduction in affinity for recognizing a monoclonal neutralizing antibody, B38. This result is significant to understand the increased transmissibility and immune invasion of two rapidly emerging SARS-CoV2 VOCs (B.1.135 and P.1). Also, the data is fundamental to assess the efficacy of different vaccines based on the spike protein.

## Acknowledgment

The author gratefully acknowledges Microsoft AI for Health program for providing Azure cloud capabilities to carry out the research (Microsoft AI for Health COVID-19 grant ID: 00011000243).

